# Comprehensive characterization of the complex *lola* locus reveals a novel role in the octopaminergic pathway via Tyramine beta-hydroxylase activation

**DOI:** 10.1101/132027

**Authors:** Nadja Dinges, Violeta Morin, Nastasja Kreim, Tony D. Southall, Jean-Yves Roignant

## Abstract

*longitudinals lacking (lola)* is among the most complex genes in *Drosophila melanogaster*, encoding up to twenty protein isoforms and acting as a key transcription factor in axonal pathfinding and neural reprogramming. Most of previous studies employed loss-of-function alleles disrupting common exons of *lola*, making it difficult to delineate its functions. To address this issue we have generated specific mutations in each isoform using the CRISPR/Cas9 system. Our targeted screen allows us to revisit the previously demonstrated roles for few isoforms and to demonstrate a specific function for one variant in axon guidance via activation of the microtubule-associated factor Futsch. Importantly, we also reveal a critical role for a second variant in preventing neurodegeneration via the control of the octopaminergic pathway. This variant is expressed almost exclusively in the octopaminergic cells and is involved in the transcriptional activation of a key enzyme of the pathway. Thus, our comprehensive study greatly expands the functional repertoire of Lola functions, and adds novel insights into the transcriptional regulatory control of neurotransmitter expression *in vivo.*

## Introduction

Size comparison of the human genome with the genome of lower organisms such as *C. elegans* or *D. melanogaster* predicts that the complexity of higher organisms does not simply rely on gene number [1]. Additional regulatory layers such as RNA editing, usage of multiple transcription start and termination sites as well as alternative pre-mRNA splicing (AS) are essential post-transcriptional mechanisms involved in the control of gene expression [2]. AS increases the number of proteins generated from a single pre-mRNA and is therefore one of the most important mechanisms to expand protein diversity. However, while the existence of cell-type specific splicing patterns is well documented there is still relatively little knowledge about the contribution of specific isoforms during differentiation or in the functionality of a given cell.

*lola* is among the most complex loci in *Drosophila* giving rise to at least 80 different mRNA isoforms through alternative *cis*- and *trans*-splicing as well as via multiple promoter activity (Figure 1A) [3–6]. In total, *lola* encodes 20 known protein isoforms (Lola A – Lola T) that contain a constitutive N-terminal BTB domain, with 17 isoforms encoding a unique zinc finger motif in their C-terminal variable exons (Figure 1B). Lola has been shown to act as a transcription factor with regulatory roles in axon growth and guidance during embryogenesis and is also required for maintaining neurons in a differentiated state by ensuring the continued repression of neural stem cells genes in neurons of the developing brain [3, 7, 8]. In addition to its described role during nervous system development, Lola has been found to control stem cell maintenance and germ cell differentiation in the *Drosophila* testis, programmed cell death during oogenesis and gonad formation in early embryo [9–11].

**Figure 1.**
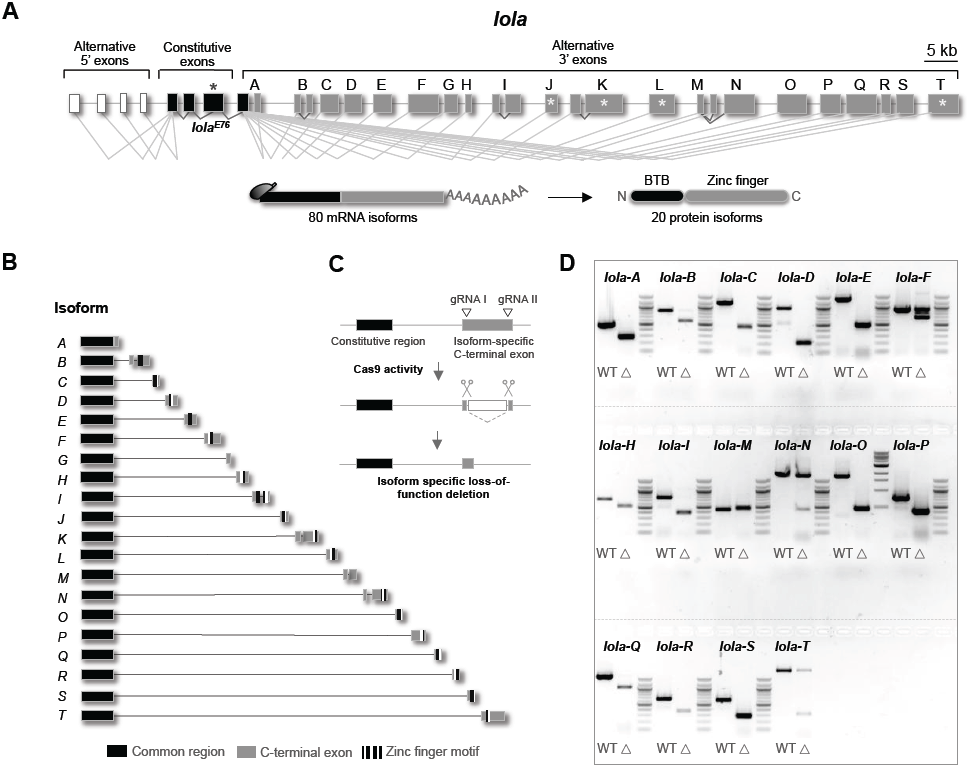
CRISPR/Cas9-induced *lola* isoform-specific knock out. See also Figures S1 and S2. **(A)** Schematic structure of the *lola* locus. *lola* comprises 32 exons including 5’ UTR exons (white boxes), constitutive exons (black) and 3’ alternative exons (grey). 80 putative transcripts encode for 20 protein isoforms, each sharing an N-terminal BTB domain but hold isoform-specific exons encoding for a C-terminal zinc-finger domain in 17 isoforms. Previously characterized mutations are marked by an asterisk. **(B)** Scheme of the 20 different *lola* isoforms (Lola-A-Lola-T), indicating sequential genomic location of C-terminal isoform-specific exons. Stripes highlight zinc-finger motifs. **(C)** CRISPR/Cas9 approach to systematically mutate each Lola isoform. Two gRNAs were designed to target the isoform-specific exon, resulting in loss-of-function deletions for respective isoforms upon Cas9 activity. **(D)** Gel analyses of established *lola* mutants. Agarose gel shows amplicons of respective genomic regions in wild type (WT, left lane) and CRISPR/Cas9 induced *lola* isoform-specific mutant flies (Δ, right lane). Mutation in Lola-M resulted in partial duplications that disrupt the frame. Mutants for *lola-G, -J, -K*, and *–L* bear a frameshift and are not displayed. Lethal alleles are heterozygous and show both the mutant and the wild type allele.

Most experiments aimed to investigate Lola functions were performed using loss of function alleles containing mutations in the N-terminal constitutive region, which affect all 20 Lola isoforms and give pleiotropic effects *in vivo* [3, 4, 7, 8, 10, 12–17]. Studies of specific isoform-mutant alleles, which are available only for Lola-K, -J, -L, and –T, revealed functions in distinct physiological processes [3, 9–11, 18]. Isoforms Lola-K and Lola-L are both involved in two unrelated mechanisms, which include motor-nerve development and germline stem cell maintenance in the male testis, indicating that at least some Lola isoforms control multiple functions during development [3, 10].

Previous studies using RNA interference (RNAi) to assess functions of individual Lola isoforms regarding neural stem cell (NSC) proliferation in the *Drosophila* brain gave contradictory results [8, 19]. Additional work demonstrated extensive and significant off-targets for RNAi experiments in *Drosophila,* underlining the importance of a permanent and specific gene knock out, in which off-targets can be evaluated [20]. Recent cutting-edge development in genome engineering now enables precise genomic manipulation by making use of the CRISPR/Cas9 technique, a system that allows to permanently alter specific DNA regions of interest [21–23].

In order to functionally characterize Lola isoforms we have performed a targeted loss-of-function screen using the CRISPR/Cas9 approach to specifically modify the *lola* locus *in vivo*. We generated loss-of-function mutations for each of the 20 known *lola* isoforms by respectively targeting the isoform-specific C-terminal exons. We found that amongst the 20 *lola* isoform-mutant strains, five are homozygous lethal during early development while three exhibit clear defects in adult flies. We confirm the previously observed lethality for the depletion of Lola-K, -L and T. Besides, we demonstrate that mutations targeting *lola-F*, in contrast to the other isoforms, result in severe disruption of axonal tracts at the ventral midline of embryos. This indicates that Lola-F is responsible for the characteristic axonal guidance phenotype observed under the complete loss of Lola. Lola-F regulates the expression of several axonal guidance genes, including the microtubule-associated factor Futsch. Remarkably, restoring Futsch expression is sufficient to rescue the axonal guidance phenotype of *lola-F* mutants, demonstrating that it is a critical target in the control of this process. Furthermore, flies bearing loss-of-function mutations for *lola-A* and *lola-H* display severe locomotion phenotypes. Finally, *lola-O* mutant flies are viable but display a strong degeneration phenotype due to a defective octopaminergic pathway. Octopamine is a critical neurotransmitter in invertebrates related to the vertebrate norepinephrine. Both octopamine and norepinephrine control diverse aspects of organismal behavior including aggressiveness, muscle activity, response to acute stress, learning and memory [24, 25]. Like other neurotransmitter their levels need to be tightly controlled. We found that Lola-O is specifically expressed in the subset of neurons that produce octopamine and regulates its biogenesis by controlling the expression of *Tyramine beta-hydroxylase* (*Tbh*), which encodes a key enzymatic component of this pathway. Together, our data provides a comprehensive functional characterization of Lola isoforms, revealing novel roles for previously uncharacterized isoforms, including an unexpected function for one variant in neurotransmitter biogenesis.

## Results

### CRISPR/Cas9-induced *lola* isoform-specific knock-out

In order to comprehensively characterize Lola isoforms *in vivo* we sought to systematically generate knock out (KO) flies for each isoform using the CRISPR/Cas9 system. Briefly, two distinct guide RNAs (gRNAs) were designed to target the isoform-specific C-terminal exon of each *lola* isoform (Figure 1C). After establishment of transgenic lines expressing each pair of gRNAs, these flies were crossed with flies expressing Cas9 in the germline. Specific modifications in the DNA of F1 flies were screened by PCR using primers spanning the flanking sequences targeted by the gRNAs, and followed by sequencing of the amplified DNA. In most cases, Cas9 activity resulted in the production of two double-strand breaks, leading to deletion of the sequence between the two Cas9 target sites and resulting in a loss-of-function mutation for the respective *lola* isoform (Figures 1D and Figure S1).

Our targeted screen resulted in the production of KO flies for all 20 known Lola isoforms. We found that mutations in 8 of these isoforms display a clear phenotype (Table 1). Five isoform-specific mutations, including in the already described *lola-K*, *-L*, and *T* as well as in *lola-F* and *-N*, result in lethality during embryonic and early larval stages. Mutations in *lola-A* and -*H* produce viable flies but they have impaired locomotion in however opposite directions (Figure S2A). While the disruption of *lola-H* leads to reduced locomotion, *lola-A* mutant flies are hyperactive. Finally, *lola-O* mutants survive until adulthood but rapidly degenerate and die around two weeks after hatching. With the exception of Lola-H and -A, we found that only the isoforms that are conserved between *D. melanogaster* and *Anopheles gambiae* play important roles during development and in adults (Table 1), raising the question of the relevance of the non-conserved Lola isoforms.

Among them, Lola-B was previously shown to play a critical role in NSC differentiation using RNA interference [19]. Upon depletion of Lola-B, proliferation of NSC was shown to be drastically reduced. However, our *lola-B* mutant flies do not recapitulate this phenotype, suggesting that this effect was likely due to off-target activity (Figure S2B). In contrast, mutations in the conserved *lola-L* and *-K* reproduce the reported defects in the innervation of ISNb motoneurons (Figure S2C), as well as the loss of *lola-T* in germ cell migration (Figure S2D). Note that the possible functional redundancy between different isoforms might prevent us to uncover additional roles for Lola. Consistent with this, reducing the level of both Lola-L and Lola-F was recently shown to alter NSC number, while reduction of either one alone has no effect [26]. In the next sections, we focus on the role of Lola-F and Lola-O, whose complete loss of function has not been previously characterized.

### Lola-F is the main isoform required for axon guidance in the developing embryo

We generated two different *lola-F* alleles that are both lethal during late embryogenesis (Figures 1D, 2A and S1). One allele specifically lacks the zinc finger motif (*lola-F*^*znf*^), whereas the second one carries a 2-bp deletion downstream of the zinc finger domain (*lola-F*^*stop*^), leading to a frameshift and premature stop codon. We confirmed the specific depletion of Lola-F protein in both homozygous mutant embryos while the overall levels of other isoforms remain virtually unchanged (Figure 2A). Immunostaining using a Lola-F specific antibody further confirmed the absence of Lola-F protein in *lola-F*^*Stop*^ mutant embryos (Figure 2B). Moreover, expression of a *lola* genomic construct rescues the lethal effect of both mutants, confirming the specificity of our loss of function alleles (data not shown). Expression analysis by *in situ* hybridization reveals a strong enrichment of *lola-F* mRNA in the developing CNS, which is lost in *lola-F*^*znf*^ mutant embryos (Figure 2C). Further analysis by fluorescent *in situ* hybridization shows localization of *lola-F* in both NSCs and differentiated neurons, suggesting possible functions during neurogenesis. At late embryogenesis and in larvae *lola-F* RNA expression becomes mainly restricted to NSCs (Figures 2C and S3A).

**Figure 2.**
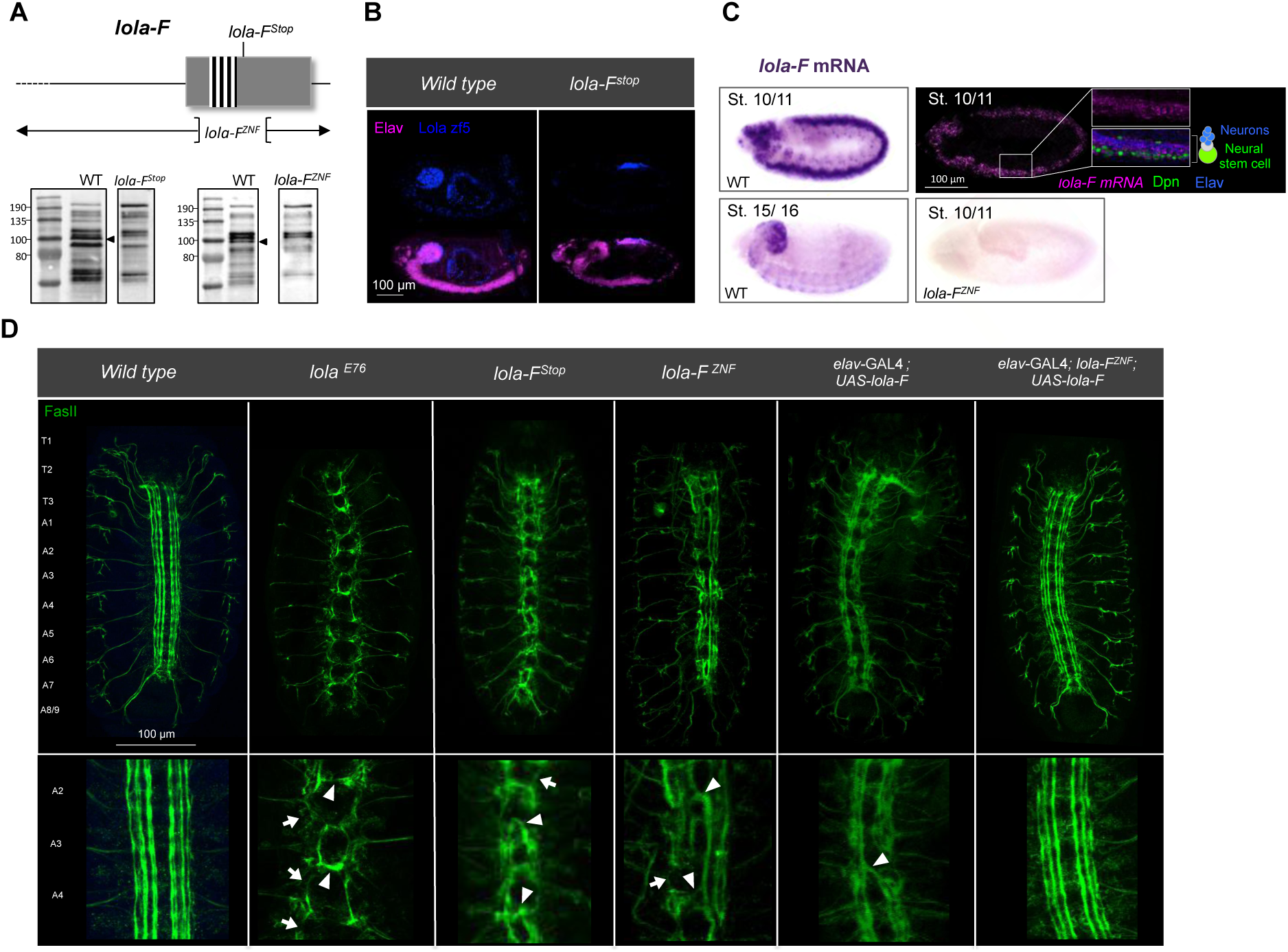
*lola-F* mutant embryos display disrupted axonal tracts. See also Figures S2 and S3. **(A)** (Top) Schematic of *lola-F* specific genomic region*. lola-F*^*Stop*^ bears a 2-bp deletion resulting in a premature stop codon downstream of the zinc-finger domain. *lola-F*^*ZNF*^ lacks the entire zinc-finger motif. (Bottom) Immunoblotting performed using an anti-Lola antibody. Arrowheads depict band specific for Lola-F. **(B)** Immunostaining using the Lola-F specific antibody Lola zf5. **(C)** *In situ*-hybridization using *lola-F* probe. *lola-F* mRNA (magenta) is enriched in the nervous system and absent in *lola-F*^*ZNF*^ mutant embryos. *lola-F* mRNA co-localizes with both the neuronal marker Elav and neural stem cell marker Dpn (lateral view, stage 10/11). **(D)** Immunostaining of the VNC using an anti-Fasciclin 2 antibody (green). Axonal pathfinding is disrupted in *lola-F* mutant embryos, similar to the defects observed for *lola*^E76^ null mutants. *lola-F* mutants display strong midline crossing of axons (arrowheads), while *lola*^E76^ null mutants show more severe axon growth defects (arrows). Neuronal *lola-F* expression in wild type disrupts axonal tracts but is sufficient to rescue the phenotype in a *lola-F* mutant background (stage 15/16 embryos, ventral view).

To address the role of Lola-F, we stained control and *lola-F* mutant embryos with anti-Fasciclin II to label axon tracts of stage 15 embryos (Figure 2D). The loss of all Lola isoforms was previously shown to strongly alter axonal growth and pathfinding at the ventral midline, a phenotype that we could recapitulate using the null *lola*^E76^ mutant allele. Remarkably, depletion of Lola-F also results in severe disruption of the ventral nerve cord (VNC). Compared to the null mutant axons extend a bit further but several crossing defects were observed. The number of neurons, NSCs or glial cells is, however, unaffected (Figures S3B and S3C). Both alleles as well as transheterozygous combinations give indistinguishable phenotypes (Figure 2D and data not shown). Moreover, expression of a *lola*-BAC was sufficient to rescue the axonal guidance defects (data not shown), confirming the specific role of this isoform in this process. We further generated a transgenic line expressing Lola-F under the control of UAS promoter. Strikingly, neuronal expression of this construct in wild type embryos strongly disrupts the VNC, mimicking the *lola-F* loss of function phenotype (Figure 2D). However, ubiquitous or neural expression of Lola-F cDNA in *lola-F* mutant embryos completely restores the axonal guidance defects (Figure 2D and data not shown). Altogether these data indicate that Lola-F plays a major role in establishing proper axonal guidance at the ventral midline and suggest that its physiological levels must be tightly controlled to ensure correct function.

We next wondered whether other isoforms that are essential during embryonic development were also involved in controlling axon pathfinding at the midline. Lola-K and –L were already shown to control muscle innervation by ISNb motoneurons. We could recapitulate this phenotype and also demonstrate that Lola-F plays a similar function (Figure S2C). However, neither the depletion of Lola-K and –L nor the absence of Lola-N and –T alter axon guidance at the ventral midline (Figure S2E). Only *lola-K* mutants display occasionally crossing defects between the A5 and A6 hemisegments. Hence, our data indicate that Lola-F is the major isoform required in axonal pathfinding at the ventral midline for which *lola* is named for.

### Lola-F activates expression of several genes involved in axon guidance

Previous studies reported the involvement of Lola in axon guidance in part by upregulating the levels of the repulsive ligand Slit and its receptor Robo at the ventral midline [15], while its effect on axon growth could be partly explained by downregulation of the actin nucleation factor Spire [16]. To get further insights into the mechanisms by which Lola controls axon growth and guidance, we took advantage of our specific *lola-F* allele to perform a transcriptome analysis of *lola-F* mutant embryos at stage 15, when majority of axon growth and guidance events are taking place. We also included samples from *lola* null mutant embryos to compare the affected genes with the absence of *lola-F*. We found that 465 genes and 586 genes were significantly up-, and down-regulated in *lola-F* mutant embryos, respectively (adjusted P-value <0.01). Intriguingly, GO term analysis revealed enrichment for genes involved in cell adhesion and axon extension specifically for down-regulated genes, underlining the role of Lola-F in regulating axogenesis (Figure 3A and Figure S4A). 217 (37 %) down-regulated genes are mutually found in the *lola* null mutant, including the previously described targets *slit* and *robo1* (Figures 3A, 3B and Figures S4B, S4C). However, in contrast to the null allele, *lola-F* mutant shows no significant change in *spire* expression (Figure 3B and Figure S4D), indicating that other isoforms must control its level. The absence of *spire* regulation by Lola-F might account for the milder effect observed on axon growth in comparison to the complete loss of *lola* (Figure 2D). Furthermore, potential antagonistic isoform-specific Lola functions might nullify each other’s effect, thus explaining that a subset of genes affected in the *lola-F* mutant is not present in the null allele.

**Figure 3.**
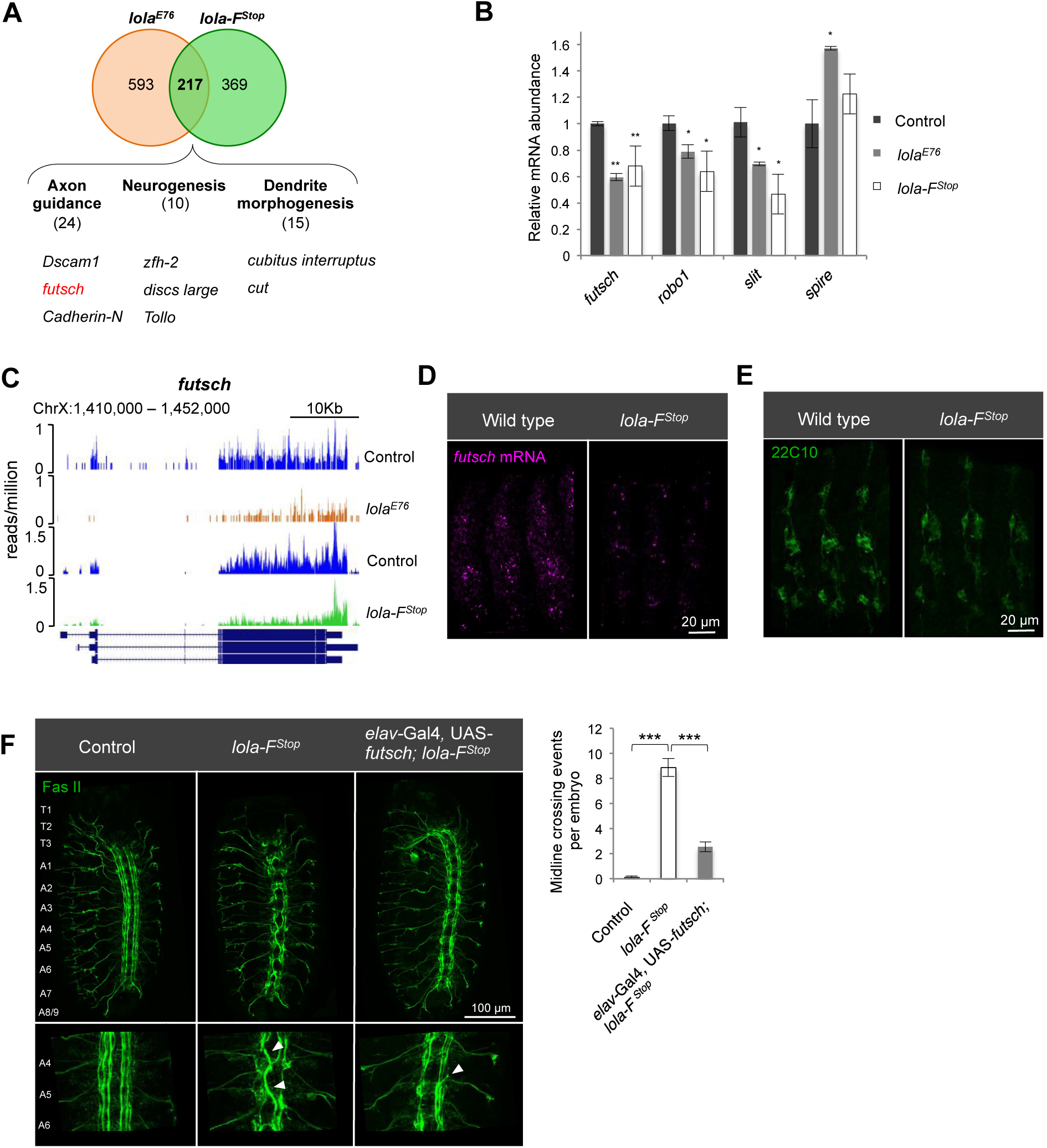
Lola-F regulates neuronal projection by activation of axon guidance genes. See also Figure S4. **(A)** Transcriptome analysis of mutant RNA extracted from *lola-F*^*Stop*^ and *lola null*^*E76*^ embryos reveals 217 commonly downregulated genes. Highlighted are examples of shared target genes involved in axon guidance, neurogenesis and dendrite morphogenesis. **(B)** qRT-PCR for selected genes on *lola*^*E76*^ and *lola-F*^*Stop*^ mutant RNA extracted from stage 15 embryos. ANOVA t-test was performed to test statistical significance. **p<0.01; *p<0.05. Data are represented as median ±SD. **(C)** Track example of poly-A selected RNA-seq at the *futsch* locus. **(D)** *In situ* hybridization to monitor *futsch* mRNA expression. *futsch* mRNA is reduced in *lola-F*^*Stop*^ mutant embryos (stage 15, lateral view). **(E)** Immunostaining using anti-22C10 antibody to monitor Futsch protein expression. *lola-F*^*Stop*^ mutants display decreased levels of Futsch protein (stage 15 embryo, lateral view). **(F**, left) Immunostaing of VNC using anti-Fasciclin II antibody to analyze midline-crossing events. Ectopic neuronal Futsch expression partially rescues axonal midline crossing of *lola-F*^*stop*^ embryos. **(F**, right) Quantification of midline-crossing events. Nine embryos were analyzed and tested for significance using ANOVA one-way t-test. ***p-value<0.001. Data are represented as average ±SEM.

Among the top 15 down-regulated genes that are common in both *lola* null and *lola-F* mutants is *futsch*, a well characterized and conserved axon guidance gene (Figure 3A). The axonal defects we observed for *lola-F* mutant embryos mimics the previously characterized *futsch*^*K68*^ mutant phenotype, as embryos show disrupted VNC motoraxons as well as stalling of the peripheral ISNb motornerve (Hummel et al., 2000, Figure S2C). The reduction in transcript levels was confirmed by qRT-PCR and *in situ* hybridization (Figures 3C, 3D and 3E). Furthermore, immunostaining on *lola-F* KO embryos using a specific anti-Futsch antibody showed an overall reduced fluorescent intensity, indicating that the protein level was also decreased (Figure 3E). To examine whether the effect on *futsch* expression in the absence of Lola-F activity contributes to the observed phenotypes, we wondered whether restoring its levels could rescue some of the *lola* defects. To this purpose, we used an EP element insertion line containing UAS regulatory elements located in the promoter region of *futsch* that we crossed with the neuronal *Elav*-Gal4 driver line (see methods). We found that embryonic progeny derived from this cross display ectopic Futsch levels (Figure S4E). Similar elevated Futsch levels were observed in embryos deficient for Lola-F. Remarkably, this ectopic Futsch expression was sufficient to partially restore both VNC and ISNb axonal defects (Figure 3F and Figure S4F). Taken together, our findings demonstrate that Lola-F regulates axonal pathfinding by activating the expression of numerous axon guidance genes during embryogenesis including the microtubule associated encoded gene *futsch*.

### *lola-O* mutant flies display a severe degeneration phenotype

We next investigated the function of Lola-O *in vivo*. We generated one mutant allele that is expected to disrupt the entire zinc finger domain (Figure 4A). The lack of mRNA expression was confirmed by qRT-PCR and RNA sequencing using RNA extracts from control and mutant strains (Figures S5A and S5B). Depletion of *lola-O* results in homozygous viable animals; yet adult flies display several abnormalities, including reduced lifespan and severe locomotion defects (Figures S5C and S5D). In addition, females exhibit cuticle malformation and suffer from partial sterility (Figures S5E and S5F). Both males and females also frequently form melanotic masses (also called pseudotumors), which occur predominantly on abdomen and limbs (Figures 4B and 4C). Melanotic masses are also occasionally observed in third instar larvae (Figure S5G). Importantly, all these phenotypes can be rescued by a genomic construct restoring *lola-O*, ruling out off-target effects. Likewise, ubiquitous or ectopic neuronal *lola-O* expression is sufficient to restore the lifespan and reduce phenotypic penetrance of *lola-O* depleted flies (Figures 4D, 4E, 4G and Figure S5H). In contrast, glial expression has no effect (Figure 4F). Taken together, these findings indicate a neuronal role for Lola-O in regulating multiple physiological functions.

**Figure 4.**
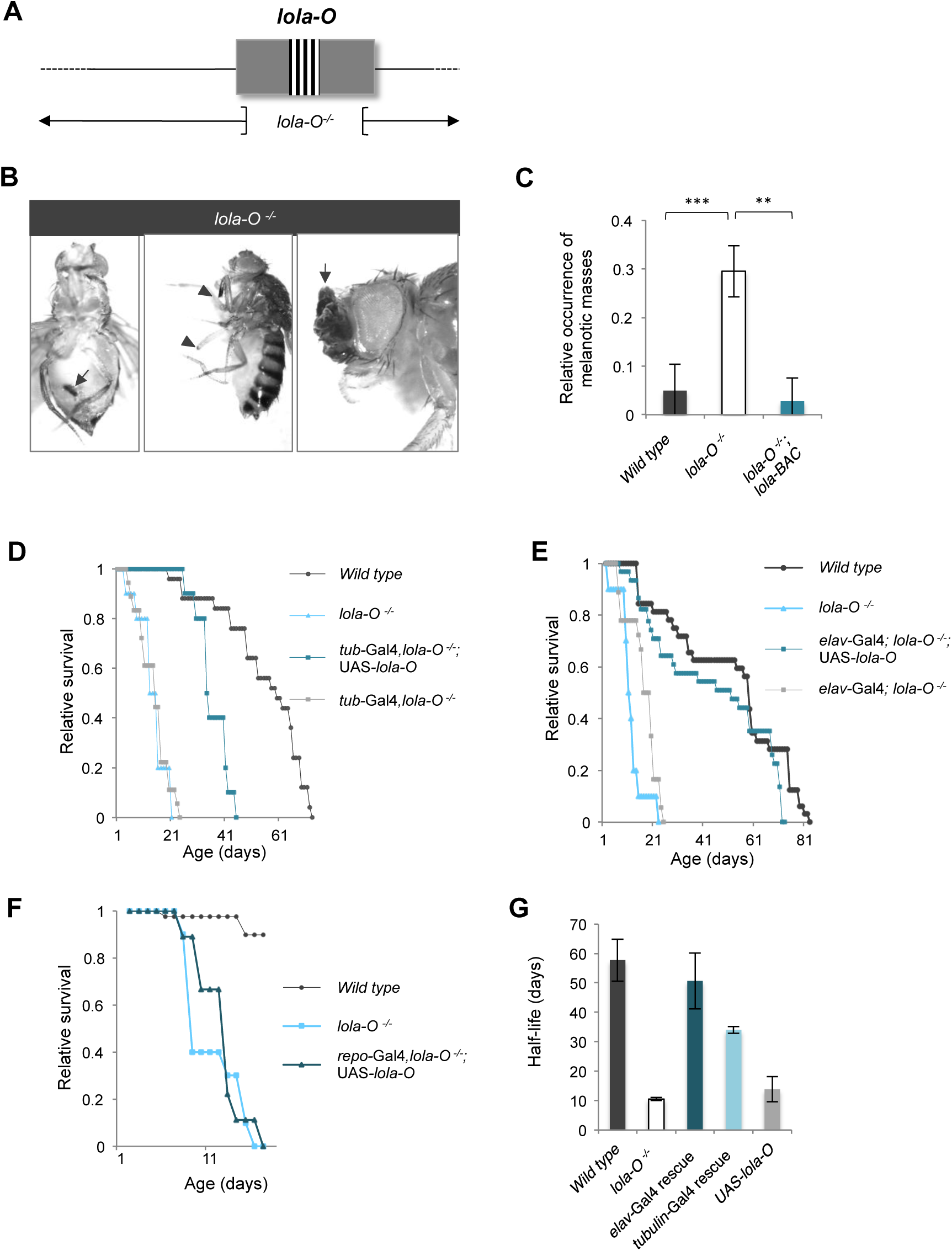
*lola-O* mutant flies display a severe neuronal degeneration phenotype. See also Figure S5. **(A)** Scheme of the *lola*-*O* genomic region and the deletion. The entire Lola-O specific zinc finger motif is deleted. **(B)** *lola-O* depleted adult flies. Lack of Lola-O induces melanotic masses (arrows), eventually leading to the loss of affected limbs (arrowhead). **(C)** Quantification of melanotic masses. Phenotypic penetrance of melanotic mass formation is reduced to control levels by recombination with a *lola* genomic construct. 10 flies were examined in four replicates at 11 days of age. Statistical analysis was performed using ANOVA one-way t-test. ***p-value <0.001.. Data are represented as average ±SEM. **(D-F)** Survival curves of adult *Drosophila*. **(D)** Ubiquitous ectopic expression of *lola-O* cDNA using *tubulin*-Gal4 elongates the lifespan of *lola-O* mutants by twofold. *lola-O* mutants recombined with the driver line serve as a control. **(E)** Neuronal *lola-O* expression using *elav*-Gal4 restores longevity of *lola-O* mutant flies to wild type levels. *lola-O* mutants recombined with the driver line serve as a control. **(F)** *repo*-Gal4 driven glial cell specific Lola-O expression is not sufficient to rescue premature lethality of Lola-O depleted flies. **(G)** Half-life quantification. Half-life is defined as the day with 50 % survival. Four biological replicates were analyzed.

### *lola-O* is specifically expressed in octopaminergic neurons

In order to obtain insights into Lola-O function, we seek to address its localization *in vivo*. For this purpose we took advantage of a fly line carrying a *lola*-BAC encoding a Lola-O-GFP fusion protein (Spokony & White, 2012). We found that Lola-O is generally expressed at a very low level in the larval brain, just at the limit of its detection. However, the low Lola-O-GFP expression is refined to only a subset of cells, which includes the midline and lateral midline of the ventral ganglion and few groups of cells in the central brain (Figure 5A). In order to reveal the identity of these cells we crossed flies carrying a UAS-GFP transgene reporter with flies expressing GAL4 under the control of various promoters, which are active in the brain. The resulting GFP expression was subsequently compared to the expression pattern of Lola-O. Interestingly, GFP expression driven by the *Tdc2*-GAL4 driver was reminiscent to the expression of Lola-O-GFP (Figure 5B).

**Figure 5.**
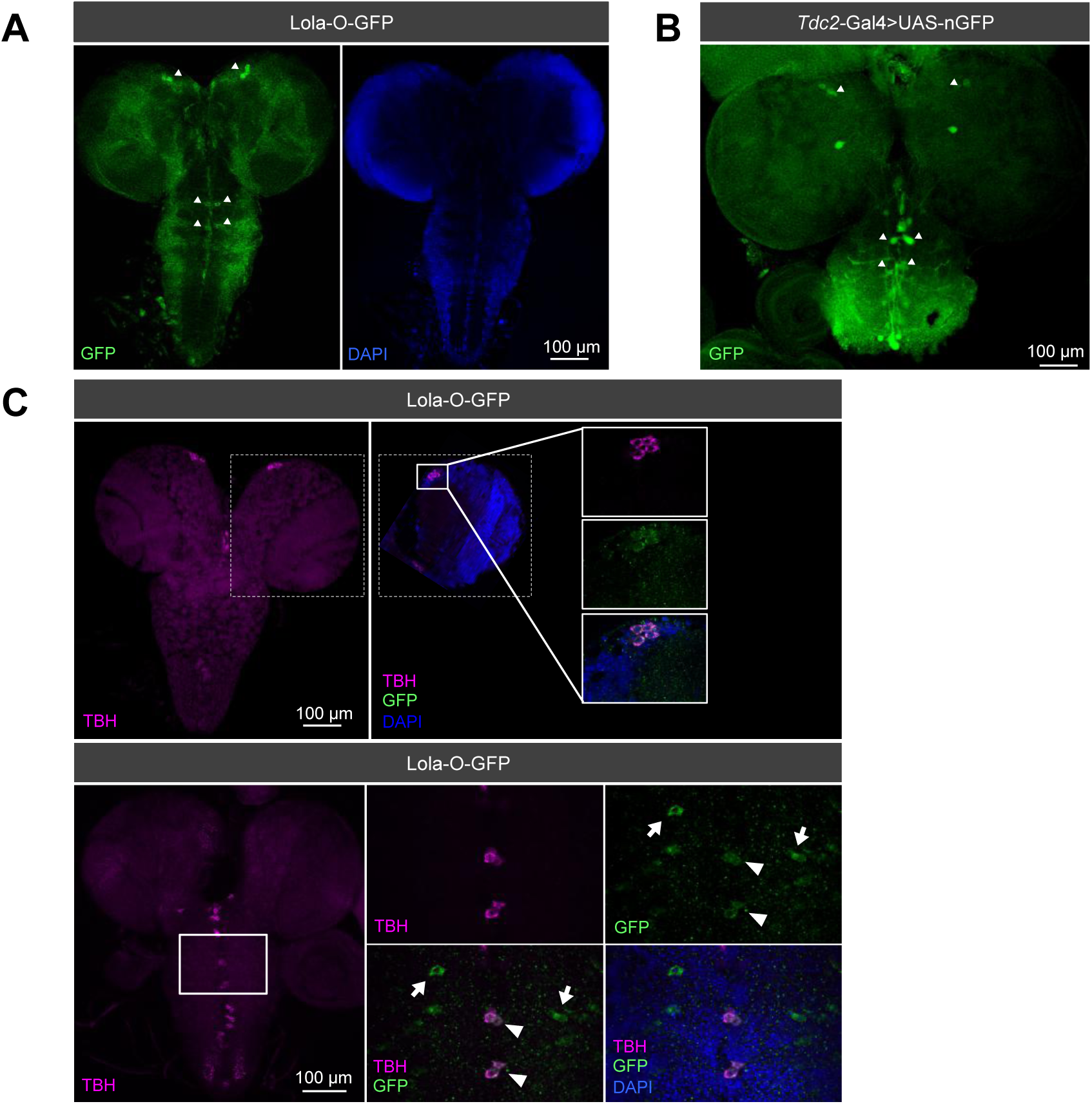
Lola-O co-localizes with octopaminergic neurons in the larval brain. **(A)** Immunostaining of Lola-O-GFP using an anti-GFP antibody (green; DAPI, blue). **(B)** *Tdc2*-Gal4 expression pattern in the larval brain revealed with an anti-GFP antibody (green). Octopaminergic cells are present in the VNC along the midline and in a subset of cells in the central brain (arrowheads), similar to Lola-O-GFP expression pattern. **(C)** Lola-O and TBH co-localize in the larval central brain and in cells along the midline of the VNC (arrowheads). Individual Lola-O positive cells at the lateral midline show no TBH immunoreactivity (arrows).

*Tdc2* stands for *Tyrosine-decarboxylase* and encodes for an enzyme required for the synthesis of octopamine (Figures 5B, 6A). Octopamine acts as a neurotransmitter, neuromodulator and neurohormone in insects and is involved in diverse physiological functions. Interestingly, perturbation of its level results in phenotypes reminiscent to the loss of Lola-O, including locomotion defects, cuticle deformity, appearance of pseudotumors and female sterility [27–29]. Its synthesis requires the amino acid tyrosine, which is modified to the intermediate compound tyramine by the enzyme Tyrosine-decarboxylase 2 (Tdc2). Subsequently, Tyramine β-hydroxylase (TBH) hydrolyzes tyramine to synthesize the neurotransmitter octopamine (Figure 6A and [30]. Noticeably, we found that mutant flies for *Tbh* (*Tbh*^*nM18*^) displayed impaired longevity, implying a previously uncharacterized role for octopamine on survival (Figure S6).

**Figure 6.**
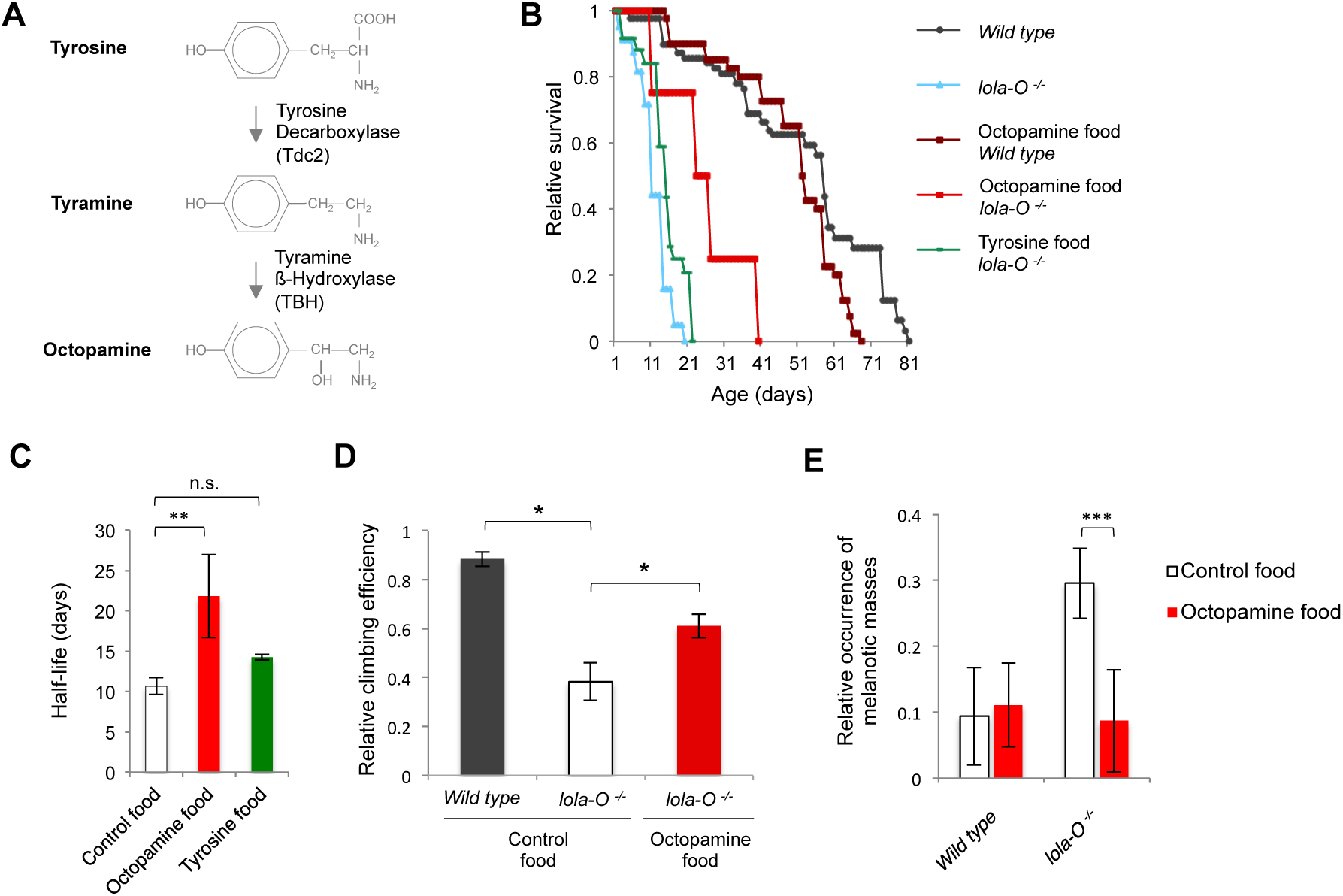
*lola-O* mutant flies can be rescued by feeding octopamine. See also Figure S6. **(A)** Scheme of the octopaminergic pathway. **(B)** Drug feeding assay for *lola-O* KO flies. Rearing *lola-O* mutant flies on octopamine-enriched food is sufficient to elongate longevity by twofold while tyrosine feeding has no profound effect on survival. The average of three biological replicates is shown. **(C)** Quantification of the half-life deduced from (B). Feeding octopamine to *lola-O* mutants increases the half-life from 10.6 days to 21.8 days. ANOVA t-test was applied, **p-value<0.001. Data are represented as average of three biological replicates ±SD. **(D)** Relative climbing efficiency. Feeding octopamine to *lola-O* mutants enhances locomotion abilities. Three days old male flies were used for locomotion quantification. ANOVA one-way t-test was performed to test statistical significance, *p-value<0.05. Data are represented as average ±SD. **(E)** Occurrence of melanotic masses. Freshly hatched males and females were separated and analyzed for melanotic masses at 11 days of age. ANOVA one-way t-test was applied, ***p-value <0.0001. Data are represented as average of four biological replicates ±SEM.

To confirm the expression of Lola-O in octopaminergic neurons, larval brains were stained for both Lola-O-GFP and an antibody that recognizes TBH as TBH expression was shown to be a faithful marker of octopaminergic neurons [31]. We found that TBH and Lola-O-GFP co-localize in a cluster of dorso-medial cells in the larval brain and along the midline of the VNC. Single cells lateral of the midline are however only positive for Lola-O-GFP (Figure 5C). Therefore, our data suggest that Lola-O function may be linked to the octopaminergic pathway.

### Expression of *lola-O* cDNA in octopaminergic neurons is sufficient to rescue most of *lola-O* mutant defects

To address a potential role of Lola-O in the octopaminergic pathway, we performed a drug-feeding assay in which *lola-O* mutant flies were supplied with octopamine-enriched food. Interestingly, ectopic feeding of octopamine to *lola-O* mutants was sufficient to elongate the half-life by twofold (12 days +/- 2.6 versus 21 days +/- 3.6, p-value <0.5), while feeding the same flies with tyrosine had no effect (Figures 6B and 6C). We verified that feeding *Tßh*^*nM18*^ mutant flies with octopamine rescues the half-life to a similar extent (Figure S6). Furthermore, octopamine-enriched food improves locomotion of *lola-O* mutants and reduces the occurrence of pseudotumors close to wild type levels (9,86% on octopamine-enriched food compared to 85,71 % on control food) (Figures 6D and 6E). Therefore, these results strongly suggest that the defects observed in *lola-O* mutant flies result from reduced levels of octopamine.

We reasoned that if Lola-O function was solely required in octopaminergic cells, expressing Lola-O specifically in these cells should be sufficient to rescue all defects associated with its loss. To test this hypothesis, we used the *Tdc2*-GAL4 driver, which is specific for octopaminergic cells. Crossing this driver line with UAS-Lola-O flies completely restores the survival of *lola-O* mutants as well as their climbing ability and the appearance of pseudotumors (Figures 7A, S7A and S7B). Furthermore, the fertility of mutant flies was also partially rescued as evidenced by the improved egg-laying rate (Figure S7C). Altogether these experiments demonstrate that Lola-O primary activity is restricted to octopaminergic neurons.

**Figure 7.**
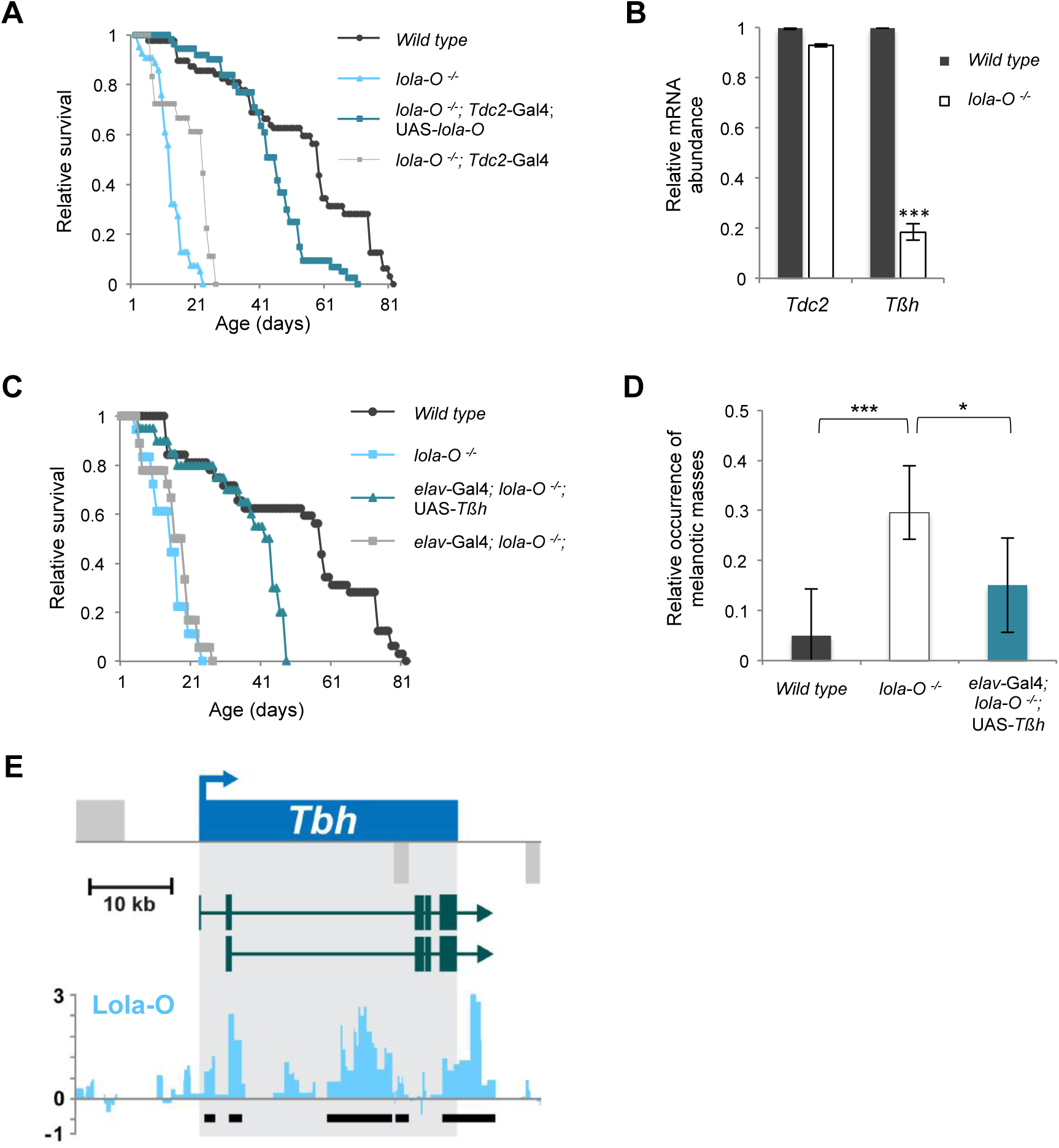
Lola-O regulates the octopaminergic pathway via the activation of *Tbh* expression. See also Figure S7. Survival curve. Expression of Lola-O in octopaminergic neurons is sufficient to restore wildtypic longevity. The average of three biological replicates is shown. **(B)** qRT-PCR using mutant RNA extract isolated from freshly eclosed adult male heads. ANOVA one-way t-test was performed. *** p-value<0.001, *p-value<0.05. Data are represented as median ±SD. **(C,D).** Neuronal *Tbh* expression using *elav*-Gal4 is sufficient to restore longevity **(C)** and occurrence of melanotic masses **(D)** of *lola-O* mutant flies. The average of two biological replicates is shown in (C). **(E)** Lola-O binding at the *Tbh* locus. Vertical bars show log2fold enrichment. Black bars highlight significantly enriched peaks.

### Lola-O regulates the octopaminergic pathway via the control of *Tbh* expression

Octopamine is synthetized via the activity of two enzymes, Tdc2 and TBH. To test whether Lola-O might control their expression *in vivo*, we analyzed transcript levels from RNA isolated from adult heads of control and *lola-O* mutant flies using qRT-PCR. Intriguingly, while *Tdc2* expression is unaffected in *lola-O* mutants, the level of *Tbh* mRNA is significantly reduced (Figure 7B). A similar result was observed in a transheterozygous combination in which *lola-O* mutant was crossed with a *lola* deficiency line (data not shown). We also found that TBH protein level is decreased as shown by reduced TBH immunoreactivity in brains of *lola-O* mutant larvae (Figure S7D). The specificity of the TBH staining was confirmed, as the signal was essentially absent in brains of *Tbh* mutant larvae (Figure S7E). Collectively, these results indicate that Lola-O is required to maintain proper *Tbh* levels.

Given the reduced abundance of *Tbh* in *lola-O* mutants we wondered whether ectopic expression of *Tbh* cDNA would be sufficient to restore longevity and phenotypic penetrance of these flies. We generated flies with integrated *Tbh* under the control of a UAS-promoter and induced expression using different GAL4-lines. Remarkably, expression of *Tbh* using a neuronal specific driver line was sufficient to rescue fly survival and to significantly decrease the occurrence of pseudotumors (Figures 7C, 7D). These findings therefore demonstrate that *Tbh* is the primary target of Lola-O in the octopaminergic pathway.

To address whether *Tbh* might be directly regulated by Lola-O we aimed to identify Lola-O genome-wide binding sites. For this purpose we took advantage of the recently established targeted DamID (TaDa) approach, which allows identifying direct target genes in a cell-type specific manner [32]. We cloned *lola-O* coding sequence into *pUASTattB-LT3-NDam* to allow the targeted, low-level, expression of N-terminally tagged Dam-Lola-O. We then induced neuronal Dam-Lola-O expression in embryos using the driver line *elav*-Gal4 and processed them at stage 17, just before larval hatching [33]. The TaDa experiment was performed in biological duplicates, showing a high degree of correlation between the two samples (r = 0.94). Consistent with *Tbh* being a direct target of Lola-O, we found that Lola-O binds *Tbh* at 5′, intronic and 3′ regions (Figure 7E), with the most significant event having a false discovery rate (FDR) of 2 × 10^−5^. Despite the specific function of this factor, Lola-O binds widely throughout the genome, with the potential to regulate about half of all genes (9158 genes identified with a FDR of < 0.01). This may reflect the potential for Lola-O to regulate different sets of genes in different tissues and/or its ability to heterodimerise with other Lola isoforms.

## Discussion

Here, we describe for the first time a comprehensive functional characterization of the different Lola isoforms *in vivo*. Using the CRISPR/Cas9 approach we generated mutants for every *lola* isoform and demonstrated that mutation in *lola-F* mimics the characteristic *lola* null mutant phenotype during embryogenesis. We further uncovered an unexpected function for Lola in the octopaminergic pathway mediated by Lola-O activity via the activation of *Tbh*. In addition to revealing novel Lola functions, this study demonstrates that the recently developed CRISPR/Cas9 system can be used to systematically address isoform-specific functions *in vivo.*

### Assigning Lola function to specific isoforms

Lola is amongst the most complex loci in *Drosophila*, encoding for 20 different protein isoforms via the usage of 3’ alternative exons. Its complete loss of function gives rise to pleiotropic defects *in vivo*, which have been difficult to analyze at the molecular level due to the paucity of specific mutant isoforms. First described in 1993 by Giniger and colleagues, Lola was shown to control axon growth and guidance in both the CNS and PNS of the *Drosophila* embryo. However, it remained unclear how it exerts these functions, and whether these effects depend on a specific Lola isoform or on the activities of multiple ones. Our results clearly establish Lola-F as being the main isoform required in these early developmental processes. A previous transcriptomic analysis from *lola*-null mutant extracts revealed only small changes on gene expression, suggesting that Lola controls axonal guidance by fine-tuning the expression of many genes involved in this process, and it is the sum of small changes on many genes that give rise to the severe *lola* null mutant phenotype [16]. Nevertheless, several key targets could be identified. For instance, in the CNS, Lola was suggested to repel longitudinal axons away from the midline through increasing the expression of the midline repellant Slit and its axonal receptor Robo [15]. In the PNS, Lola controls ISNb axonal growth partially via reducing the expression of the actin nucleation factor Spire [16]. In spite of Lola-F being involved in both processes, we found that only *slit* and *robo* expression was altered, while *spire* was unchanged. Furthermore, our study revealed *futsch* as being a key target of Lola-F in axon growth. Futsch is a microtubule-associated protein whose expression levels need to be tightly controlled. For instance, heterozygous animals display mutant phenotypes, including slower growth rates and motor system abnormalities, indicating that its dosage is critical for its function [34]. Moreover, its expression in the somatic and visceral musculature is repressed by Tramtrack [35]. It is interesting to note that Tramtrack, like Lola, encodes for a zinc finger and BTB-domain containing protein, suggesting that this class of proteins plays a prominent role in either repressing or activating Futsch expression. Previous studies on several BTB-containing proteins demonstrated that the BTB domain mediates homomeric and heteromeric dimerization, and transcriptional repression through the recruitment of diverse co-repressor proteins [36, 37]. Our data, however, indicates that Lola-F is a general activator of neuronal genes. Further interactome studies will therefore be necessary to understand how different BTB proteins exert antagonistic molecular functions on gene expression.

Interestingly, the expression of Lola-F drops as *Drosophila* development progresses and becomes primarily restricted to NSCs. This finding suggests that neuronal gene expression outside the NSCs must be maintained by different means, such as via the activity of other transcription factors or via an epigenetic mechanism. Intriguingly, Lola-F was shown to interact early in development with the histone H3S10 kinase JIl-1 [6]. Via this molecular activity, JIL-1 maintains euchromatic regions by antagonizing Su(var)3-9-mediated heterochromatinization. Therefore, it is possible that early during development, Lola-F establishes an epigenetic state compatible with gene expression via its association with JIL-1 and once established, its function might become dispensable. The remaining Lola-F expression in NSCs appears important to maintain their differentiation capacity as the double KD of Lola-F and Lola-L leads to dramatic overprofileration of NSCs in the central brain [26]. It will be important to further identify the direct targets of Lola-F and Lola-L in this process to understand how these two isoforms cooperate to allow NSC differentiation.

### Novel connection between Lola and the octopaminergic pathway

Our study also revealed a novel role for Lola in the octopaminergic pathway. This function was probably missed due to pleitotropic effects of *lola* null mutants combined with the lack of specific *lola-O* alleles. Our findings demonstrate a regulatory role for Lola-O in the octopaminergic pathway by activating *Tbh* expression, which encodes an enzyme required for the synthesis of octopamine, a monoamine that acts as a neurohormone neuromodulator as well as neurotransmitter [38]. Monoamine neurotransmitter levels are usually tightly regulated, as their misregulation can lead to a wide range of disorders in human [39]. Accordingly, several evidences indicate that the absolute levels of octopamine must also be tightly controlled. For instance, reducing its levels in a *Tbh* mutant leads to diverse defects, including female sterility, locomotion, aggressiveness and pseudotumor formation [24, 27, 28]. On the other hand, increasing its level also leads to similar abnormalities [29]. Despite this dosage-function dependency, very little is known about the mechanisms controlling octopamine synthesis during different stages of the *Drosophila* life cycle. Studies on honeybees have shown that octopamine synthesis increases with age [40], which might be also the case in *Drosophila*. Additionally, octopamine synthesis has been shown to be stress-induced in the haemolymph in both locusts and cockroaches [41], implying that differential concentrations of octopamine are required in response to altered environmental circumstances. Recent reports also demonstrated an increase in octopamine metabolism upon heat conditions in *Drosophila*, which is mediated via FOXO activity [42]. It would therefore be interesting to test whether TBH expression is also stress-induced and whether Lola-O contributes to this effect.

### Future Directions

Our studies illuminate a novel role for Lola in controlling neurotransmitter signaling. Interestingly, Lola-O expression appears almost restricted to TBH positive cells in the larval nervous system, albeit at very low levels. The mechanisms that restrict Lola-O expression to this subset of cells are currently unknown. It is widely known that promoter activity can influence and determine tissue-dependent expression of individual genes, suggesting that expression of Lola-O in octopaminergic neurons might be a consequence of alternative promoter activity. As *lola* is encoded by at least four promoters, it will be interesting to determine their impact on the expression levels of the different isoforms. Alternatively, a splicing regulator might be specifically expressed in octopamine responsive cells, promoting the expression of Lola-O exclusively in these cells. Additional experiments are needed to further reveal the mechanism of the restricted expression of Lola-O in the brain and to address whether analogous mechanisms apply in vertebrates to control the level of norepinephrine upon normal and disease conditions.

## Author contributions

Conceptualization, J.Y.R and N.D.; Methodology, J.Y.R., N.D. and V.M.; Investigation, N.D. and V.M.; Computation analysis of RNA-Seq, N. K.; Computational analysis of DamID-Seq, T.D.S; Writing – Original Draft, N.D.; Writing –Review & Editing, J.Y.R. and N. D.; Funding Acquisition, J.Y.R.; Supervision, J.Y.R.

## Acknowledgments

We would like to thank the Bloomington Drosophila Stock Center for fly reagents; The Developmental Studies Hybridoma Bank for antibodies; the Drosophila genomics Resource Center for plasmids, M. Monastirioti for Tbh alleles and TBH antibody; E. Giniger for Lola antibody; C. Berger for Deadpan antibody; members of the Roignant lab for fruitful discussion; the IMB Genomics, Bioinformatics and Microscopy Core facilities for tremendous support; and A. Dold, C. Berger and J. Urban for critical reading of the manuscript. Research in the laboratory of J.-Y.R. is supported by the Marie Curie CIG334288 and the Deutsche Forschungsgemeinschaft (DFG) SPP1935 Grant RO 4681/4-1.

## Accession number

RNA-Seq and DamID-Seq data have been submitted to GEO under accession number GSE97836.

Reviewer link: https://www.ncbi.nlm.nih.gov/geo/query/acc.cgi?token=ixexgyyavrgxdwf&acc=GSE97836

## EXPERIMENTAL MODEL AND SUBJECT DETAILS

### Fly stocks

*Drosophila melanogaster w*^*1118*^ was used as wild type control. For rescue experiments the following driver lines were used, *Tdc2-*GAL4*, tub*-GAL4, and *elav*^*C155*^-GAL4*. Df(2R)ED2076* served for generation of *trans*-heterozygous *lola* mutants. *PBac(lola.J-GFP.FLAG)VK00033* and *PBac(lola.GR-GFP.FLAG)* were used for rescue experiments (Spokony & White, 2012). For transcriptome analysis, *lola*^*E76*^ and *lola-F*^*Stop*^ flies were balanced with *w*; P{sqh-mCherry.M}* to enable selection of homozygous embryos based on fluorescence (Martin *et al*., 2009). *P{w[+mC]=EP}futsch[EP1419]* served for the rescue experiment of *lola-F*^*Stop*^ embryos. Flies were obtained at the Bloomington *Drosophila* Stock Center. *Tbh*^*nm18*^ mutant flies were a kind gift from M. Monastirioti

### CRISPR/Cas9 mutant flies

gRNA sequences were cloned into pBFv-U6.2B [43], sequenced and injected (250 ng/μl) in our lab into *y1 v1 P(nos-phiC31\int.NLS)X; attP40.* Transgenic flies were further crossed with *y*^*2*^ *cho*^*2*^ *v*^*1*^*; attP40(nos-Cas9)/CyO* and flies from the F1 generation were PCR screened for the expected deletion mutation using primer sequences flanking the gRNA sequences. Obtained PCR amplicons were purified and sequenced at GATC Biotech.

### UAS constructs

Coding sequences of *lola-O, lola-F* and *Tbh* was amplified from cDNA using Phusion High Fidelity Polymerase (NEB) and inserted into Gateway plasmids with N-terminal Flag-Myc or Flag-HA tag (pPFMW or pPFHW, respectively; obtained from *Drosophila* Genomics Resource Center at Indiana University). All constructs were sequenced prior to injection into *w*^*1118*^. *Drosophila* germline injection for *lola-F* and *Tbh* was performed in-house by standard procedure. The construct for UAS-*lola-O* was injected at BestGeneInc.

### Drug Feeding Assay

Mutant flies of the respective strain were collected within 10 hours of eclosion, gender separated and placed on medium containing 5 mg/ml or 7.5 mg/ml octopamine or tyrosine for males and females, respectively. Flies were examined daily for survival rate and phenotypic penetrance. 20 flies were used for each condition in three individual biological replicates. Octopamine was additionally fed to *w*^*1118*^ flies.

### Locomotion assay

20 freshly hatched male and female flies of the respective mutant stock were separated and either directly placed into measuring cylinders or staged until desired age. The locomotion was assessed using the climbing assay described previously [44]. Flies were tapped to the bottom and flies passing 8 cm in 10 or 5 seconds, respectively, were counted. Measurements were repeated five times in three independent biological replicates.

### Lifespan assay

20 control or experimental flies were collected within 10 hours of eclosion and maintained on standard medium at 25°C. Survival was analyzed every two days and flies were transferred to new vials twice a week. Experiments were performed in three biological replicates.

### Fertility assay

10 female virgin flies were separated upon hatching and mated with 5 males for three days. Upon fertilization, flies were transferred onto fresh agar plates every twelve hours. Number of eggs laid was counted for five days. Experiments were performed in three biological replicates.

### Transcriptome analysis

Collection of *lola* mutant embryos: Embryos were collected at 25°C for two hours and subsequently developed for 13 hours. *lola-F*^*FS*^ and *lola*^*E76*^ embryos were hand-sorted and collected based on the expression of mCherry. Control embryos were collected in parallel and both mutant and control embryos were subsequently transferred into TRIzol reagent (Thermo Fisher Scientific) and RNA was isolated using the manufacturers protocol.

Collection *of lola-O* mutant adult flies: 20 males of either *lola-O* mutant or control flies were collected within 10 hours of eclosion, transferred into TRIzol reagent and subjected to RNA isolation.

### cDNA library preparation

RNA was isolated, DNase I (NEB) treated according to the manufacturers protocol and subjected to library preparation using the NEBNext® Ultra™ Directional RNA Library Prep Kit for Illumina®. 1 μg of total RNA was used as starting material for library preparation. cDNA libraries were subsequently submitted for high throughput sequencing on a HiSeq2500.

### Computational analysis

Libraries for transcriptome analysis were sequenced in two different sequencing runs on a HiSeq2500. *Lola-F*^*stop*^ and control samples were sequenced paired end. *lola*^*E76*^ and control were sequenced single read. Demultiplexing and fastq conversion was done with bcl2fastq (v. 1.8.4). Reads were mapped using STAR (v. 2.5.0c) against ensembl release 79 (BDGP6). For *lola-F* knock out and the corresponding control the first read was used for mapping to ensure comparability with *lola* Null and corresponding control samples. Mapped reads were filtered for rRNA and mitochondrial RNA reads before further processing.

Counts per gene were calculated using htseq-count (v. 0.6.1p1) with ensembl release 79 as a reference. Differentially expression analysis was done using DESeq2 (v. 1.10.0) with an FDR filter of 1%.

### GO Term method and Plot outline

GO Terms overrepresentation was calculated using GOStats (v. 2.38.1) requiring a minimal amount of 5 genes per GO Term and adjusted *p*-value smaller than 1%. Afterwards, terms were summarised using semantic similarity (GOSemSim 1.30.3). Only GO Terms with a OddsRatio larger then 5 are displayed.

### Targeted DamID

*UAS-Dam-lola-O* flies were generated by amplifying the full coding sequences and cloning it into pUASTattB-LT3-NDam (kind gift from A. Brand). Upon microinjection and generation of transgenic flies, *UAS-LT3-Dam-lola-O* flies were crossed with *elav*-Gal4 to induce neuronal expression of Dam-Lola-O. Analysis of Lola-O binding sites was performed on stage 17 embryos (20-22 hours AEL) for *UAS-LT3-Dam* (control) and *UAS-Dam-LT3-lola-O* flies. Genomic DNA isolation and subsequent treatments were performed as described (Marshall et al., 2016). Purified and processed genomic DNA of two biological duplicates was subjected to library preparation using the NebNext DNA Ultra II library kit (New England Biolabs) and sequenced on a NextSeq500. The first read was mapped to *Drosophila melanogaster* genome (BDGP6) using bowtie (v. 2.2.9), binned to GATC fragments and normalized against the Dam-only control (Marshall and Brand, 2016). Peaks were called and mapped to genes using a custom Perl program (available on request). In brief, a false discovery rate (FDR) was calculated for peaks (formed of two or more consecutive GATC fragments) for the individual replicates. Then each potential peak in the data was assigned a FDR. Any peaks with less than a 1% FDR were classified as significant. Significant peaks present in both replicates were used to form a final peak file. Any gene within 5 kb of a peak (with no other genes in between) was identified as a potential target gene.

### qRT-PCR

Total RNA was prepared and transcribed into cDNA using MMLV reverse transcriptase (Promega). For the measurement of RNA levels qRT-PCR analysis was performed using a ViiA7 real-time PCR system (Applied Biosystems). Measurements were done in triplicates. Relative RNA levels were normalized to *rpl15* levels. Primer sequences are listed in Supplemental Tables.

### *In situ* hybridization

Preparation of digoxigenin-labeled RNA probes: primers were designed to amplify a unique region within *lola* exons with the reverse primer containing the SP6 sequence. The PCR was performed on embryonic cDNA using Phusion DNA Polymerase (NEB). 250 ng of template PCR product was used to perform in vitro transcription using SP6 RNA polymerase (Roche) and DIG labeled UTP (Roche). The reaction was incubated over night at 37°C, EtOH precipitated and resuspended in DEPC water to obtain a concentration of 100 ng/μl. Probes were diluted 1:50 in hybridization buffer for in situ hybridization. Embryos were fixed for 25 minutes in fixation solution (400 μl PBS, 500 μl n-heptane, 130 μl 37% formaldehyde) while shaking at RT. After washing in MeOH several times embryos were gradually transferred into PBTween (PBS; 0.3 % Tween20), followed by three washes for 15 minutes, and finally into HB4 hybridization buffer (50 ml formamide, 25 ml 20xSSC, 200 μl Heparin (50 mg/ ml), 100 μl Tween20, 500 mg Torula Yeast RNA extract). After equilibration at RT, embryos were pre-hybridized in HB4 at 56°C for several hours. Upon denaturation of the diluted RNA probe at 80°C for 10 minutes, embryos were hybridized over night at 65°C. Embryos were subsequently incubated in washing buffer (formamide: 2xSSC, 0.1 % Tween20; 1:1) for 30 minutes at 65°C and transferred into PBTween at RT before they were incubated with anti-DIG-AP antibody (1:1000, Roche) over night at 4°C. Upon several washes in PBTween and one rinse with AP buffer (100 mM NaCl, 50 mM MgCl2, 100 mM Tris pH 9.5, 0.1 % Tween20) probes were visualized using the NBT/BCIP solution in AP buffer (1:00). For fluorescent *in situ*-hybridization, embryos were incubated with anti-DIG-HRP (1:1000, Roche) and probes were visualized using the tyramide signal amplification (Alexa Fluor 568, Thermo Fischer).

### Whole mount embryo immunostaining

Embryos were dechorionated in 50% bleach for 2 minutes and fixed for 25 minutes in 500 μl n-heptane, 400 μl 1xPBS, 130 μl 37% Formaldehyde. Wash solution was PBS with 0.3 % Triton X-100. Primary antibodies used were mouse anti-Fas II (1:20; 1D4, DSHB), rat anti-Elav (1:100; 7E8A10, DSHB), mouse anti-Repo (1:100; 8D12, DSHB), mouse anti-22C10 (1:50, DSHB), mouse anti-Lola zf5 (1:100, 1D5, DSHB), guinea pig anti-Deadpan (1:50, kind gift from C. Berger), rat anti-TBH (1:75, kind gift from M. Monastirioti), rabbit anti-GFP (1:500; TP401, Torrey Pines Biolabs). Appropriate combinations of secondary antibodies (Jackson Immunoresearch Laboratory) were applied. Samples were analyzed with a Leica SP5 confocal microscope.

### Larval L3 brain immunostaining

Larval CNS was dissected in cold PBS, fixed for 20 minutes in 4% Paraformaldehyde in PBS and subsequently treated as embryonic samples. Mounting was done in Vectashield and a confocal stack was recorded using Leica SP5 confocal microscope.

### Western blotting

For western blot analysis, staged embryos of the corresponding genotype were collected and homogenized in lysis buffer (140 mM NaCl, 10 mM Tris-HCl pH 8, 1 mM EDTA pH 8, 0.5% Triton X-100). The protein extract was separated on 8% SDS-PAGE gel and western blot analysis with affinity purified anti-Lola antibody (1:500, kindly provided by E. Giniger) was performed by standard methods. For visualization, ultra-sensitive enhanced chemiluminescent reagent (Thermo) was used.

## QUANTIFICATION AND STASTISTICAL ANALYSIS

Statistical parameters and significance are reported in the Figures and the Figure legends. For comparisons of the means of two groups, student’s t-test was used. For comparisons among more than two groups the one-way ANOVA was performed followed by multiple comparisons using t-tests with Bonferroni normalization of p-values.

